# A biosynthetic platform for antimalarial drug discovery

**DOI:** 10.1101/814715

**Authors:** Mark D. Wilkinson, Hung-En Lai, Paul S. Freemont, Jake Baum

## Abstract

Advances in synthetic biology have enabled production of a variety of compounds using bacteria as a vehicle for complex compound biosynthesis. Violacein, a naturally occurring indole pigment with antibiotic properties, can be biosynthetically engineered in *Escherichia coli* expressing its non-native synthesis pathway. To explore whether this synthetic biosynthesis platform could be used for drug discovery, here we have screened bacterially-derived violacein against the main causative agent of human malaria, *Plasmodium falciparum*. We show the antiparasitic activity of bacterially-derived violacein against the *P. falciparum* 3D7 laboratory reference strain as well as drug-sensitive and resistant patient isolates, confirming the potential utility of this drug as an antimalarial. We then screen a biosynthetic series of violacein derivatives against *P. falciparum* growth. The demonstrated varied activity of each derivative against asexual parasite growth points to potential for further development of violacein as an antimalarial. Towards defining its mode of action, we show that biosynthetic violacein affects the parasite actin cytoskeleton, resulting in an accumulation of actin signal that is independent of actin polymerization. This activity points to a target that modulates actin behaviour in the cell either in terms of its regulation or its folding. More broadly, our data show that bacterial synthetic biosynthesis is a suitable platform for antimalarial drug discovery with potential applications in high-throughput and cost-effective drug screening with otherwise chemically-intractable natural products.

## INTRODUCTION

Malaria has a huge global health burden, with around half of the world’s population at risk of contracting the disease which killed over 400 000 people in 2017^1^. Malaria disease is caused by apicomplexan parasites from the genus *Plasmodium*, with *Plasmodium falciparum* causing the majority of deaths worldwide. The symptoms of malaria disease develop during the asexual stages of the parasite life cycle, which occurs in the blood stream. Here, the parasite undergoes multiple rounds of growth, replication and invasion of red blood cells. Various drugs have been developed to target the asexual stages of the parasite but, inevitably, resistance has evolved to every major front-line therapy for malaria treatment including, most recently, artemisinin combined therapies (ACTs)^2^. Multi-drug resistance to ACTs, focussed in the Greater Mekong Subregion of South East Asia, has been reported both as delayed parasite clearance and, more worryingly, treatment failure^3^. The challenges of emerging drug resistance combined with the cost associated with development of new drugs make it essential to explore new ways to develop novel antimalarial compounds.

Previous work identified violacein, a violet indolocarbazole pigment produced by bacteria (Fig. 1a), as a potential antimalarial able to kill both asexual *P. falciparum* parasites *in vitro* and protect against malaria infection in a mouse malaria model *in vivo*^4–6^. Violacein’s antimalarial activity has therefore identified it as a potential for future drug development. However, commercial violacein samples can only be obtained through laborious purification from bacteria (*Chromobacterium*^7,8^ or *Janthinobacterium*^9^) because of the complexity of its highly aromatic structure (Fig. 1a). Purification from these bacteria requires specialised equipment and high-level biosafety equipment since these bacteria themselves can cause deadly infections^10^. As such, commercially available violacein is extremely expensive. Alternative strategies of violacein synthesis are being explored, in particular the use of synthetic biology to engineer industrial bacterial species that can express non-native violacein. Several groups including ours^11^ have been successful in implementing a five-gene violacein biosynthetic pathway (vioABCDE) into *Escherichia coli* or other heterologous hosts^12–14^, providing a route for robust, in-house and inexpensive compound production.

**Fig. 1:**
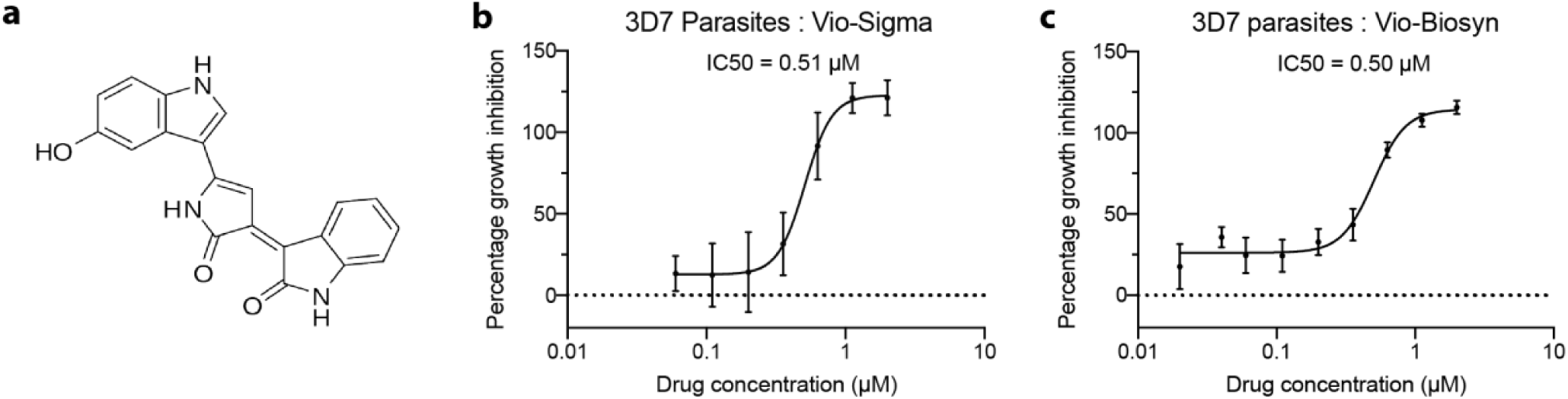
*Plasmodium falciparum* asexual growth inhibition assays with violacein. **a** The chemical structure of violacein (PubChem CID 11053) **b-c** Commercially available violacein (b) and biosynthetic violacein (c) kill asexual 3D7 parasites with a half maximum inhibitory concentration of 0.51 µM (Vio-Sigma) and 0.50 µM (Vio-Biosyn).

We have previously extended the success of this biosynthetic pathway by generating combinations of 68 new violacein and deoxyviolacein analogues. These combinations are achieved by feeding various tryptophan substrates to recombinant *E. coli* expressing the violacein biosynthetic pathway or via introduction of an *in vivo* chlorination step -the tryptophan 7-halogenase RebH from the rebeccamycin biosynthetic pathway^13,15–17^. This biosynthetic approach is able to produce large quantities of compound derivatives using simple, cheap and non-hazardous bacteria compared to native producing strains in a sustainable and flexible approach.

Here, we set out to explore whether use of this biosynthetic system could be developed as a route to antimalarial compound production and testing, measuring the activity of derivatives on the growth of *P. falciparum* sexual and asexual parasites. We have confirmed the viability of the system, ensuring there is no background antiparasitic activity in bacterial solvent extracts lacking violacein. We then tested the biosynthetic violacein extract from *E. coli* and confirmed its half maximal inhibitory concentration (IC50) in agreement with a commercial violacein standard and previous studies^14^. Finally, as well as using this approach to explore the mode of action of violacein, we show that extracts representing a diverse series of biosynthetically derived variants show varying effects on parasite growth, with 16 of the 28 compound mixtures inhibiting growth to a greater level than the parent violacein molecule. Indeed, one purified compound, 7-chloroviolacein, exhibits a ∼20% higher inhibition activity than the underivatized violacein compound. The high-throughput approach used in this study suggests that biosynthetic systems may therefore provide an, as yet, untapped resource for screening complex compounds and optimising them for antimalarial discovery.

## RESULTS

### Violacein expressed using synthetic operons kills *P. falciparum* 3D7 parasites

Previous work has shown that violacein is able to kill asexual *P. falciparum* parasites *in vitro*^9,13^ but noted erythrocytic rupture at concentrations above 10 µM and a cytotoxic IC50 of 2.5 µM. Taking this into consideration, we used concentrations of 2 µM violacein or less to explore growth inhibition of *P. falciparum* asexual stages, noting no phenotypic effect on erythrocyte morphology at the highest final concentration (Fig. S1). Our biosynthetic system for violacein production requires chloramphenicol drug pressure, which is known to affect parasite viability^18^. We first set out to ensure presence of this known antibiotic did not affect parasite growth. Extract from bacteria lacking the violacein producing enzymes but grown under chloramphenicol pressure (i.e. background) did not affect parasite viability (Fig. S2). This gave us confidence that extracts from biosynthetically modified *E. coli* would report only on the activity of a drug produced but not from background chloramphenicol contamination. To test this, we compared the activity of a commercial violacein standard (Vio-Sigma) with violacein derived from bacterial solvent extracts from *E. coli* biosynthesis (Vio-Biosyn) on wild-type 3D7 *P. falciparum* growth, using a well-established asexual growth inhibition assay. No difference in the IC50 values between the two violacein samples was seen (Figs. 1b-c). We further tested the two violacein samples on sexual parasites by measuring exflagellation^19^ and saw no difference in the IC50 values of around 1.7 µM (Fig. S3) but complete IC50 curves could not be generated without going above the cytotoxic threshold of 2.5 µM. All these data demonstrate that solvent extracted violacein from *E. coli*, Vio-Biosyn, is active and that its production provides a suitable platform for development and testing of potential antimalarial compounds.

### Biosynthetic violacein extract kills both artemisinin-resistant and -sensitive field isolate parasites

To explore whether violacein has utility for addressing emerging ACT drug resistance in the field, we tested the efficacy of Vio-Sigma and Vio-Biosyn on two parasite clinical isolates deriving from the Greater Mekong subregion, where ACT resistance is concentrated. Both clinical isolates have been phenotypically characterised in clinic, showing either treatment failure or success, adapted to culture and genotyped for the C580Y Kelch-13 resistance marker^20^ known to correlate with sensitivity to artemisinin-based drugs. Both the artemisinin-sensitive isolate (ANL1, Kelch13 wild-type) and the artemisinin-resistant isolate (APL5G, Kelch13 C580Y) were sensitive to Vio-Biosyn and Vio-Sigma with similar IC50 values (Figs. 2a-d). Activity against artemisinin-resistance provides support for development of the violacein scaffold for addressing emerging drug-resistance in the field.

**Fig. 2:**
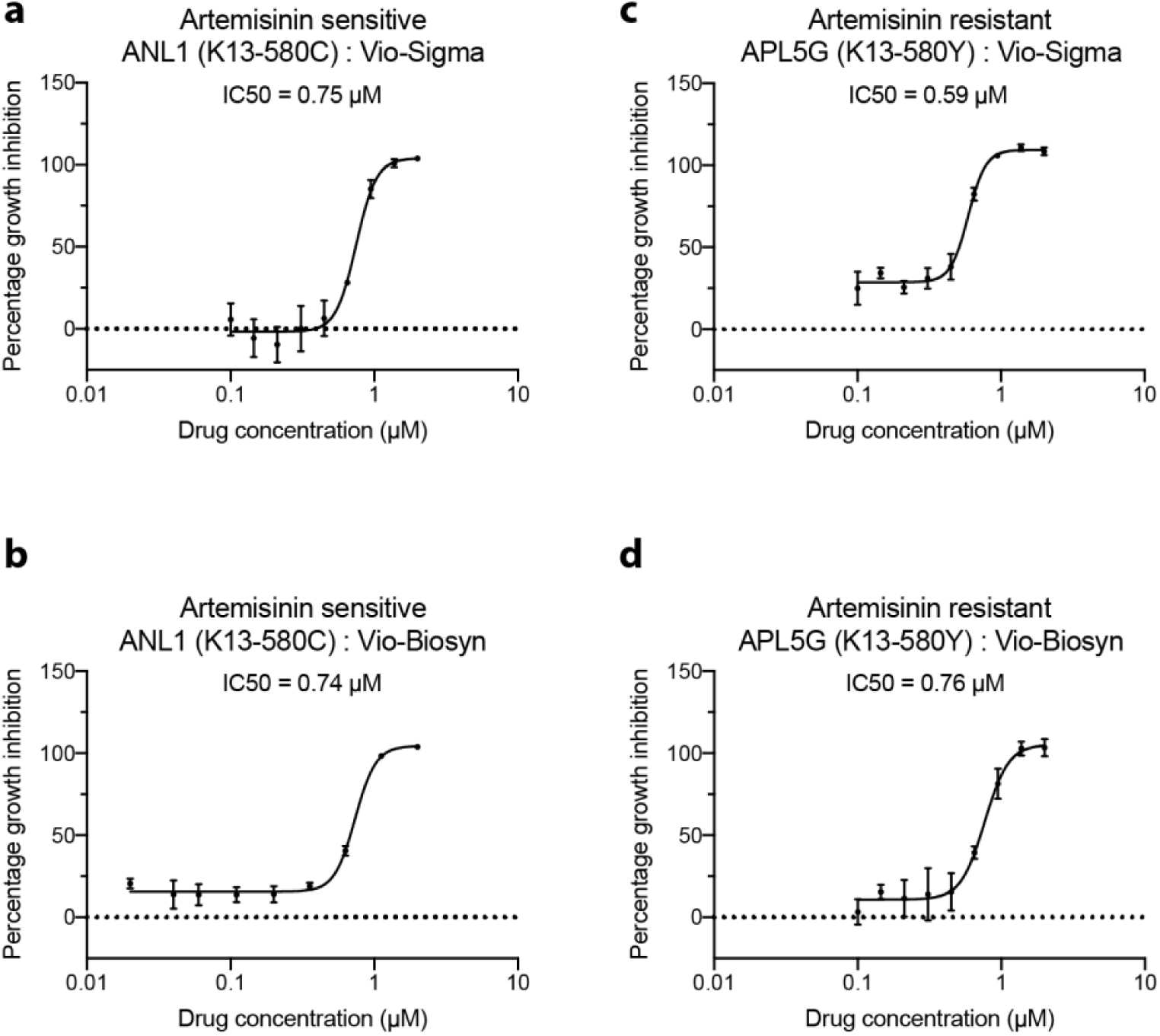
Biosynthetic violacein kills both artemisinin-sensitive and -resistant parasite clinical isolates. **a-d** Both commercially available violacein and biosynthetic violacein are able to kill parasite clinical isolates either sensitive (ANL1/Kelch13 wild-type, a,b) or resistant to artemisinin (APL5G/Kelch13 C580Y, c,d).

### Violacein derivatives show potent anti-malarial activity

To explore whether bacterial biosynthesis could be utilised further to generate compound derivatives, increasing the throughput of complex molecule testing, we obtained extracts from 28 bacteria strains each modulated to synthesise a mixture of violacein analogues (Fig. S4)^21^. The bacterial extracts were produced by feeding corresponding tryptophan substrates as described previously^14^ and violacein concentrations in the extracts were calibrated against a violacein standard. Asexual growth assays were again carried out, testing each extract at the IC50 of biosynthetic violacein, 0.50 µM. We saw a large variation of inhibition of parasite growth, with 8 compound mixtures exhibiting >95% inhibition, whilst 12 others showed a decreased effect on parasite growth (Fig. 3a). As a proof-of-concept, one of the more active extracts was used to purify the violacein derivative, 7-chloroviolacein (Fig. 3b). 7-chloroviolacein exhibited a decrease in IC50 from 0.51 µM (violacein) to 0.42 µM (7-chloroviolacein) (Fig. 3c). The speed and low cost of extracting these violacein analogues and purifying them directly from bacteria suggests an entirely new approach to complex compound drug-testing for antimalarial discovery and optimisation.

**Fig. 3:**
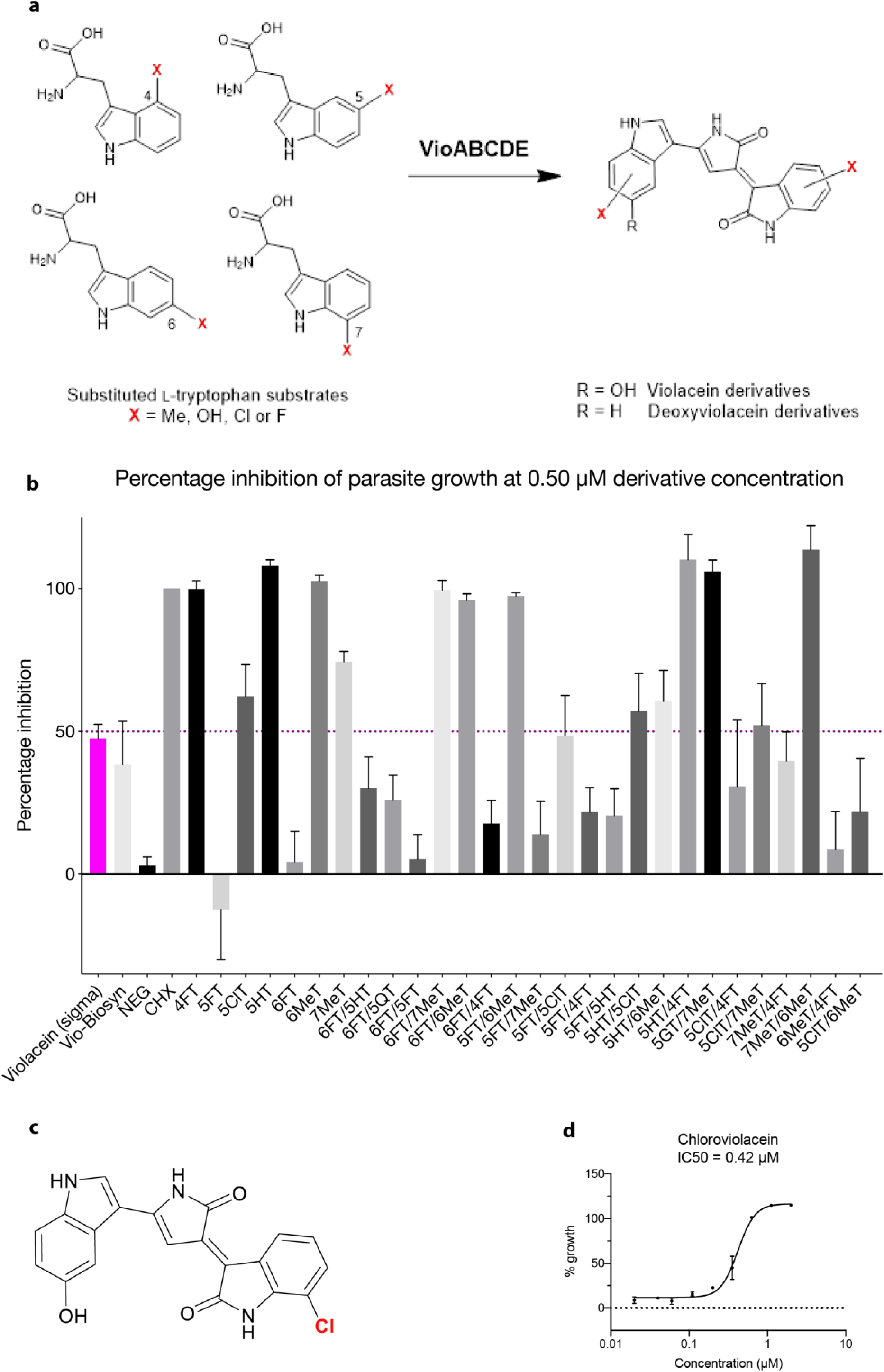
*Plasmodium falciparum* growth inhibition assays testing a series of biosynthetic violacein derivatives. **a** Tryptophan derivatives (left) used to generate violacein derivatives (right). **b** A screen of violacein derivative mixtures at 0.50 µM (adjusted by measuring absorbance at 575 nm) shows some mixtures have more potent antiparasitic ability than others. **c** 7-chloroviolacein **d** The activity of purified violacein derivative, 7-chloroviolacein, shows an IC50 value of 0.42 µM against 3D7 wild type parasites.

### Biosynthetic violacein affects actin dynamics in the cell but does not affect polymerisation *in vitro*

The mode of action of violacein against *P. falciparum* parasites has not previously been characterised. Towards exploring the phenotype associated with its activity we performed flow cytometry and immunofluorescence assays (IFAs) to observe any changes in the parasite under biosynthetic violacein treatment. A 3D7-derived parasite line expressing a constitutive cytoplasmic green fluorescent protein (sfGFP) marker was labelled with DNA marker DAPI (4′,6-diamidino-2-phenylindole) and a monoclonal antibody that preferentially recognises filamentous actin^22,23^ to explore overall parasite morphology: cytoplasm, nucleus and actin cytoskeleton respectively. Parasites were then treated with either negative, dimethyl sulfoxide (DMSO), or positive, actin filament stabilising compound, jasplakinolide (JAS) controls as well as Vio-Biosyn. Parasites were checked by flow cytometry for any differences in overall signal (Fig. S5). A low actin-positive signal was seen with DMSO treatment, as expected given the predominance of short, transient filaments and globular actin in asexual parasites^24^. The intensity of actin labelling following JAS and violacein treatment both, however, showed marked increases when compared to DMSO (Fig. S5).

To explore the nature of flow-cytometry changes in actin intensity following violacein treatment, IFAs were undertaken. In the DMSO-treated parasites, the GFP signal is spread throughout the cytoplasm along with a clearly defined nucleus as expected (Fig. 4a). The actin signal is diffuse with a low background (Fig. 4a). Following JAS treatment, actin filaments are known to be stabilised^24^ producing an expected concentrated overall actin signal (Fig. 4b), indicative of high local concentrations of polymerized actin. Parasites treated with Vio-Biosyn also gave a much higher actin signal than untreated controls, though distinct from that following JAS treatment (Fig. 4c). The concentrated signal from Vio-Biosyn was broader across the cell and less focussed in localised regions of the cell periphery. This matched the overall intensity of signal seen by flow-cytometry and relative to sfGFP signal as a control for parasite size (Fig. 4d). In the DMSO-treated control, the diffuse actin signal is 3% of total GFP signal. This increases to 27% upon JAS treatment, indicative of an increased number of filaments, whereas parasites treated with Vio-Biosyn reach a mean average signal 98%, representing a huge increase in actin accumulation in the cell. This broad concentration of actin signal would be indicative of either massively increased filament nucleation or actin aggregation as caused by actin misfolding. To test whether Vio-Biosyn directly affects actin filament formation (as JAS does), both drugs were assayed using a pyrene-labelled actin assembly assay, used previously to test compound derivatives for actin activity^24^. No effect on actin polymerisation was seen with Vio-Biosyn when compared to either JAS (filament nucleating) or the monomer-stabilising drug latrunculin B (Fig. 4d). Together, these observations suggest Vio-Biosyn does not directly interact with actin. It is therefore possible that Vio-Biosyn interacts with actin indirectly such as via the known actin-binding partners in the parasite cell^25^ or via an alternative pathway involved in actin folding, which would give rise to actin aggregation within the cell.

**Fig. 4:**
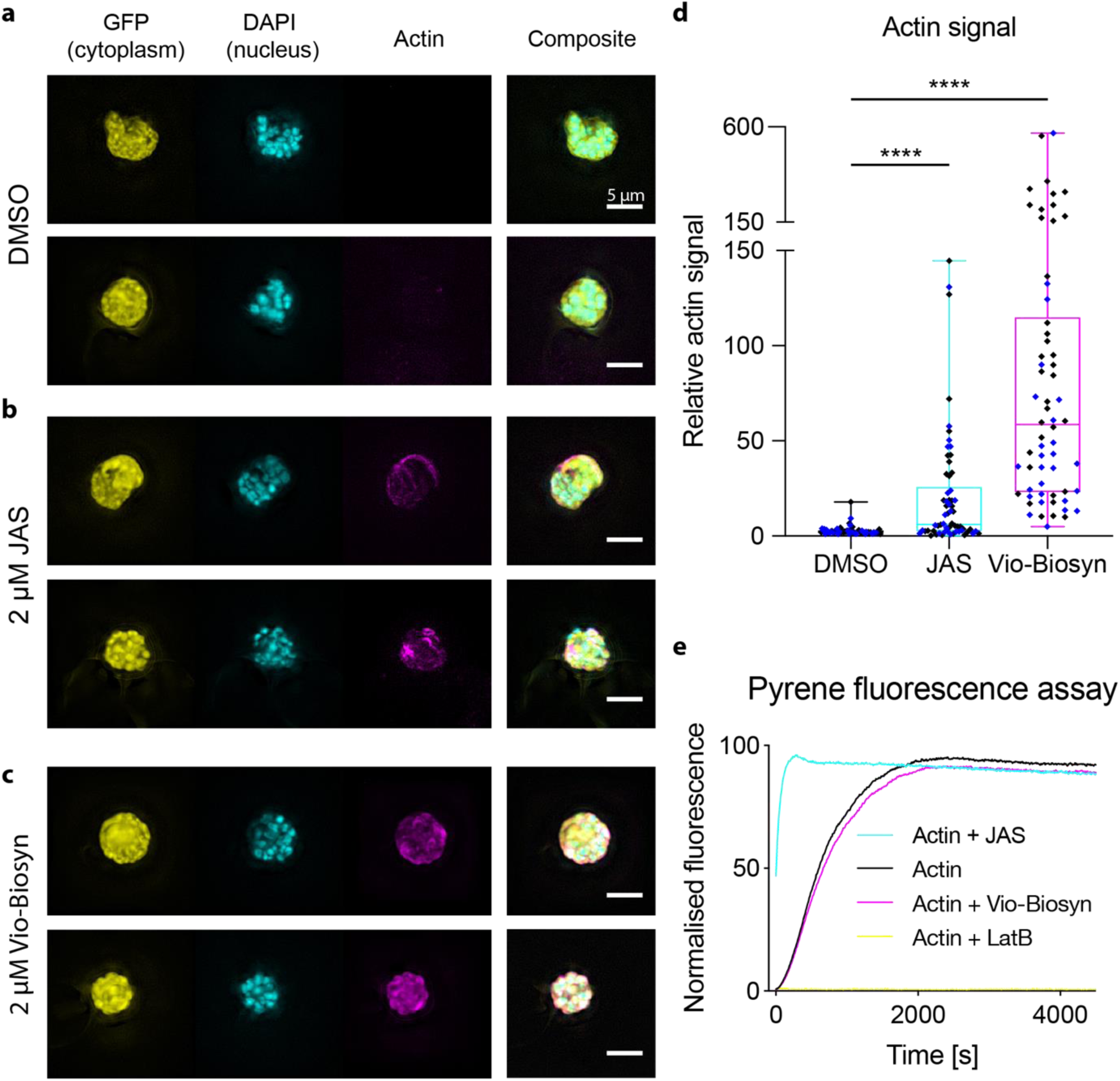
The phenotype of biosynthetic violacein treatment on cytoplasmic GFP expressing 3D7 parasites suggests modulation of the actin cytoskeleton through indirect action. **a** 3D7 parasites expressing cytoplasmic GFP, treated with DMSO have a diffuse actin signal. **b** Parasites treated with 2 µM Jasplakinolide (JAS), which stabilises filament formation, have clearly formed actin filament structures that localise to the parasite cell periphery. **c** Parasites treated with 2 µM biosynthetic violacein have increased local actin concentrations both around the outside of the cell and in the centre. **d** Parasites treated with biosynthetic violacein have a much greater overall actin signal than both DMSO control and JAS treated cells, as measured by actin fluorescence relative to cytoplasmic GFP signal. Data shown is of 62 images over two biological repeats (black and blue) for each sample. P values are unpaired students t-test, **** : p < 0.0001. **e** Biosynthetic violacein shows no effect on actin polymerisation *in vitro* as measured by pyrene-actin polymerization, compared to two known actin binders (Latrunculin B, which binds to the monomer and prevents filament formation, and Jasplakinolide, which increases nucleation and stabilises actin filaments). LatB, Latrunculin B; JAS, Jasplakinolide. Scale bar = 5 µm.

## DISCUSSION

The emergence of resistance to front-line artemisinin-based drug treatments for malaria is a major threat to global health. As such, new antimalarial treatments are in urgent demand. Here, we tested violacein, a compound with known antibacterial, antitumorigenic and antiparasitic activity, against *P. falciparum* parasites validating its potential utility for antimalarial drug development. We showed that biosynthetically produced violacein was as effective as commercially available violacein, with a mode of action that affects the actin cytoskeleton of the parasite. We also tested 28 violacein analogue mixtures using a high-throughput growth assay on asexual parasites suggesting this method of biosynthetic production is a suitable platform for antimalarial discovery and optimisation.

Previous work has shown that violacein is capable of killing lab-derived chloroquine-resistant *P. falciparum* parasites^14^. Here, we showed that patient-sourced clinical isolates, sensitive or resistant to artemisinin, could equally be killed by both commercial and biosynthetic violacein with similar IC50 values. Furthermore, although full IC50 curves could not be generated, the compounds both inhibit development of the sexual stages of the parasite life cycle with an IC50 of around 1.7 µM (Fig. S3). Any compound identified using this assay with an IC50 of less than 2.0 µM is considered for further compound development^26^. Combined, these data suggest violacein is a potential drug that could be developed to antagonize resistance in the field and target both asexual and sexual stage parasites. Whilst the derivative library tested consisted of mixtures of violacein analogues, it is encouraging to see some of these compound mixtures have considerably more potent anti-malarial activity than violacein itself. Critically, when we tested a purified compound (7-chloroviolacein) we saw a noticeable reduction in IC50 (0.51 µM to 0.42 µM), illustrating the potential of biosynthetic production of antimalarial compounds for rapid screening and rational drug optimisation.

Interestingly, violacein-treated parasites have cytoskeletal deformities that suggested disruption to actin modulation within the parasite. Given violacein has no effect on actin polymerisation kinetics *in vitro*, it is possible that the phenotype observed is as a result actin aggregation in the cell, which could be a side effect of actin misfolding. *P. falciparum* requires actin as an essential part of its motor complex and for other processes in the cell^9^. Unlike most proteins, actin requires a dedicated chaperonin system to fold into its native state^27^. Of note, this entire pathway is highly upregulated in artemisinin-resistant parasite isolates^28^ and would constitute a well-justified target for antagonising drug resistance in the field. Further work in testing the effects of violacein on actin folding or modulation are clearly required to explore whether this is the target for the drug. Ultimately, the ability of violacein to affect such a major pathway as actin dynamics in the cell, as well as kill drug-resistant parasites, provides an encouraging outlook for its therapeutic development.

In summary, our data show that a bacterial biosynthetic platform for creating compounds and their derivatives is suitable for testing for antimalarial drug development. As the need for novel therapeutics increases and interest in natural compounds, often complex in nature, grows we hope to use this approach to develop novel chemical scaffolds in a high throughput manner towards finding the next generation of antimalarials.

## METHODS

### Generation and extraction of violacein and derivatives

Violacein and derivatives were generated and extracted as previously described^29^. Briefly, *E. coli* JM109 strain (Promega) was transformed with the violacein pathway plasmid (pTU2S(VioAE)-b-VioBCD) and grown overnight before being inoculated into LB broth until OD600 reaches 0.5. These cultures were then supplemented with either tryptophan or a synthetic tryptophan analogue at 0.04% (w/v) and grown at 25 °C for up to 65 h before pelleted at 4000 rpm. The cell pellet was then resuspended with 1/10^th^ volume of ethanol to extract violacein, followed by centrifugation to separate ethanol supernatant containing violacein extract from cell debris. The supernatant was then dried *in vacuo* and stored at -20 °C for long-term storage or reconstituted in DMSO for growth inhibition assays. Concentrations of violacein in the bacterial solvent extracts were calibrated against commercial violacein standard (Sigma) based on absorbance at 575 nm. Compound mixtures used in the growth inhibition assay consist of mixtures of violacein derivatives (Fig. S4) as described previously^14^.

### *Plasmodium falciparum* growth inhibition assays

*P. falciparum* parasite lines 3D7, ANL1, APL5G were used for the GIAs. The 3D7 sfGFP line (G. Ashdown *et al*, manuscript in preparation) was used for immunofluorescence assays. All parasite lines were cultured in complete RPMI (RPMI-HEPES culture media (Sigma) supplemented with 0.05 g/L hypoxanthine, 0.025 g/L gentamicin and 5 g/L Albumax II (ThermoScientific) and maintained at 1% to 5% parasitaemia and 4% haematocrit. For the growth inhibition assay (GIA), 96-well plates were pre-dispensed with a serial dilution of compound and normalised to 1% DMSO. Double-synchronised ring-stage parasites (1% parasitaemia, 2% haematocrit, complete media, sorbitol synchronised at ring stage at 0 hrs and 48 hrs) were added to the wells to a total volume of 101 µL. Cultures were incubated for 72 h at 37 °C in a gas mixture of 90% N_2_, 5% O_2_, 5% CO_2_. Red blood cells were lysed through freeze-thaw at -20 °C and parasites were resuspended and lysed with 100 µL lysis buffer (20 mM TRIS-HCL pH 7.5, 5 mM EDTA, 0.008% saponin, 0.8% Triton-X 100) containing 0.2% SYBR green and incubated for 1 h at room temperature. SYBR green fluorescence (excitation 485 nm / emission 535 nm) was measured using a Tecan infinite M200 Pro. Data shown is the mean average of 3 biological replicates (± SEM), each of which is a mean average of 3 technical replicates (unless stated otherwise), and is normalised to a positive control (cycloheximide) and a negative control (DMSO only). IC50 values were calculated using GraphPad Prism version 8.0.

### Immunofluorescence assays

100 µL of mixed stage sfGFP parasite line (5% parasitaemia, 2% haematocrit) was incubated for 24 h with 2 µM of the compound of interest. At t=0 h and t=6 h, parasites were fixed with 4% paraformaldehyde, 2% glutaraldehyde (Electron Microscopy Sciences) and incubated on a roller for 20 mins at room temperature (RT), before being pelleted at 3000 RPM and washed three times in 100 µL 1 X phosphate-buffered saline (PBS). The cells were subsequently permeabilised in 0.1% Triton-X 100 (Sigma) for 10 mins at RT before being pelleted and washed three times in PBS as before. Cells were blocked in 3% bovine serum albumin (BSA) in PBS for 1 h at RT on a roller before being incubated with primary antibody (1:500 mouse anti-actin 5H3^14^) for 1 h at RT. Cells were washed three times with PBS before addition of the secondary antibody (1:1000 anti-mouse Alexa 647 conjugated) for 1 h at RT. Cells were washed three times in PBS and resuspended in 100 µL PBS with 0.05% DAPI. Cells were diluted 30-fold and loaded onto polylysine-coated coverslips (ibidi) before being imaged. Imaging was performed on a Nikon Ti-E microscope using a 100 x Plan Apo 1.4 NA oil objective lens with ‘DAPI’, ‘FITC’ and ‘Cy5’ specific filtersets. Image stacks were captured 3 µm either side of the focal plane with a z-step of 0.2 µm. Image analysis was conducted on raw image data sets in ImageJ, calculating a ratio between AlexaFluor 647 and FITC by measuring the mean signal intensity in a defined area of 88 nm^2^. 62 images were captured for each sample from two wells from two biological repeats. Images shown in Fig. 4 were deconvolved in Icy using the EpiDemic Plugin and a maximum intensity projection was made in ImageJ.

## List of abbreviations

DAPI: 4′,6-diamidino-2-phenylindole
IFA: immunofluorescence assay
GIA: growth inhibition assay
JAS: Jasplakinolide
LatB: Latrunculin B

## DECLARATIONS

### Availability of data and materials

The datasets used during the study are available from the corresponding author(s) upon reasonable request.

### Author contributions

MDW planned and performed the experiments and wrote the manuscript under the guidance of JB. HEL created the violacein constructs and extracted violacein and its derivatives under the guidance of PSF. All authors read, edited and approved the final manuscript.

### Ethics approval and consent to participate

Not applicable.

## Acknowledgements

Clinical isolates were provided by Professor Kesinee Chotivanich and Dr Huy Rekol (Director of the National Centre for Parasitology, Entomology and Malaria Control) through a joint Medical Research Council (MRC) and National Science Technology Development Agency (NSTDA) award (MR/N012275/1 to JB), acknowledging support from the TRAC clinical studies, funded through the UK Government Department for International Development (DFID). The 3D7 sfGFP line was provided by Dr Kathrin Witmer (Imperial College London). We wish to acknowledge Thomas Blake for his help conducting the flow cytometry and Alisje Churchyard for conducting the exflagellation inhibition assay.

## FIGURES

### VISUAL ABSTRACT

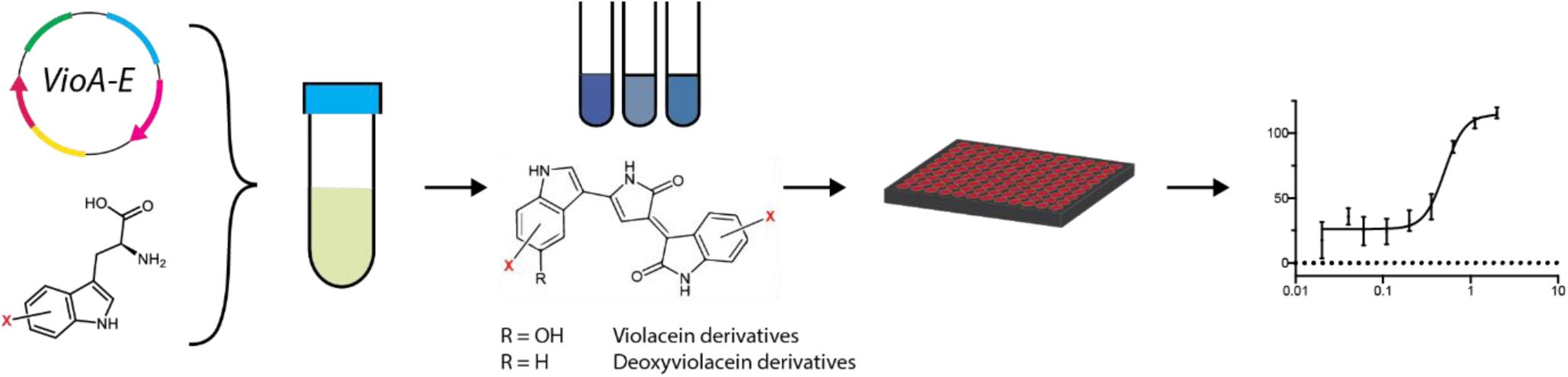

Purification of violacein and derivatives for use in the growth inhibition assay. *E. coli* cells were transfected with the violacein biosynthetic genes (*vioABCDE*) and fed with either tryptophan or its analogues as precursors for violacein analogue biosynthesis. Violacein and its derivatives were solvent extracted and serially diluted to test against various strains of *P. falciparum* parasites.

**Fig. S1:**
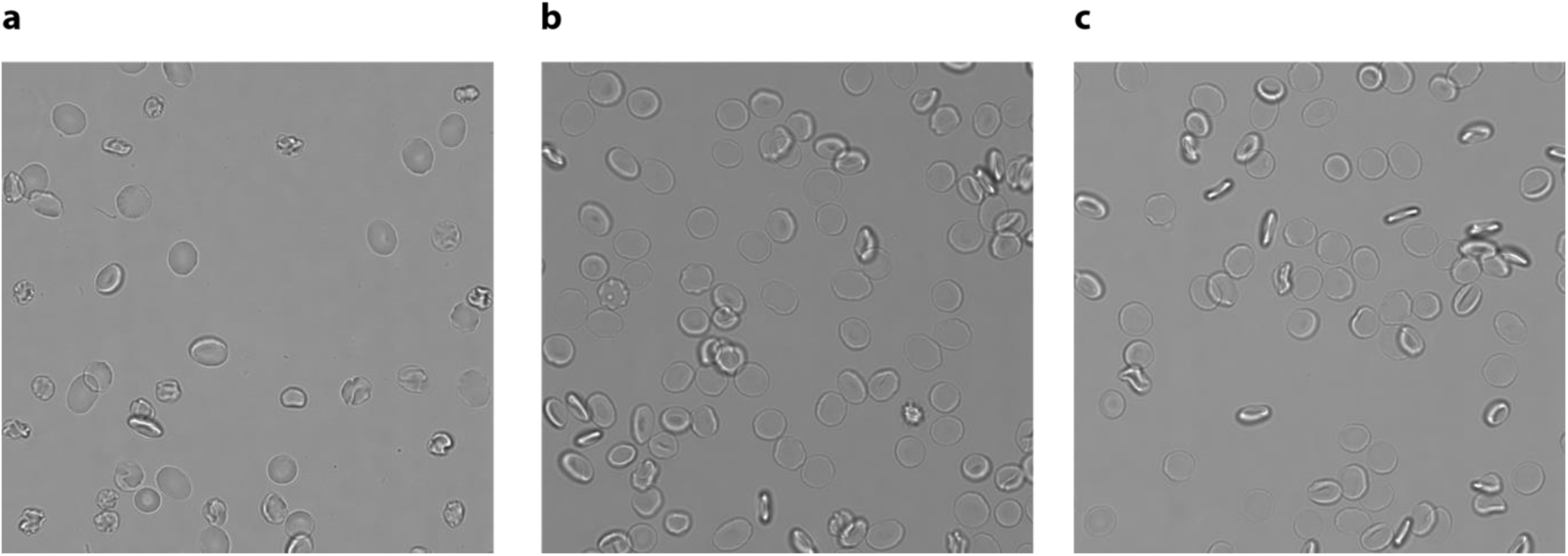
Violacein does not affect red blood cell viability the maximum concentration used in these assays. **a** DMSO-treated red blood cell control **b** Red blood cells treated with 2 µM violacein (Sigma) **c** Red blood cells treated with 2 µM biosynthetic violacein.

**Fig. S2:**
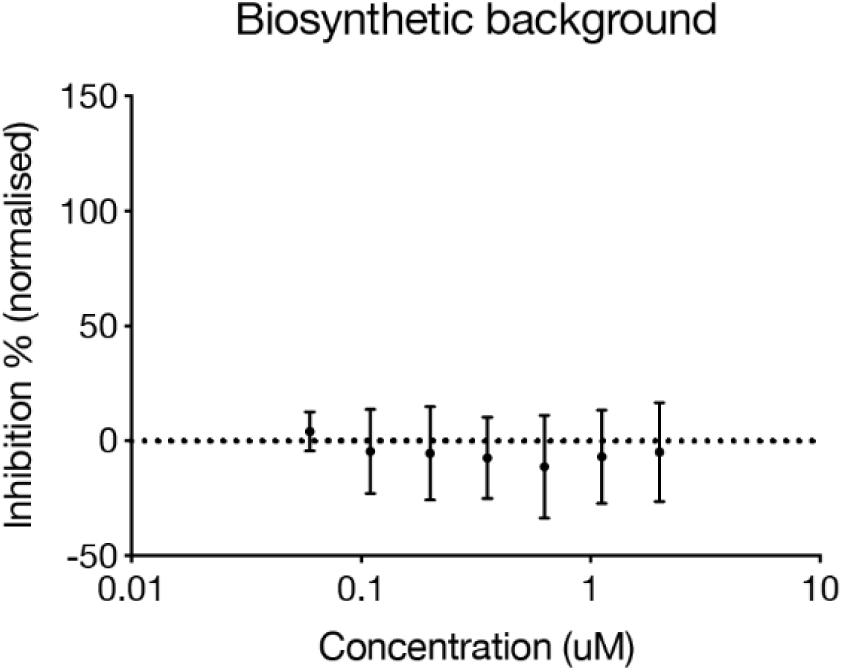
Extract of *E. coli* cells without the violacein biosynthetic genes tested against *P. falciparum* 3D7 wild type parasites. The concentration of extract was adjusted to correspond to that of violacein-containing extract by normalising to the same wet cell mass. Data shown is of three technical replicates.

**Fig. S3:**
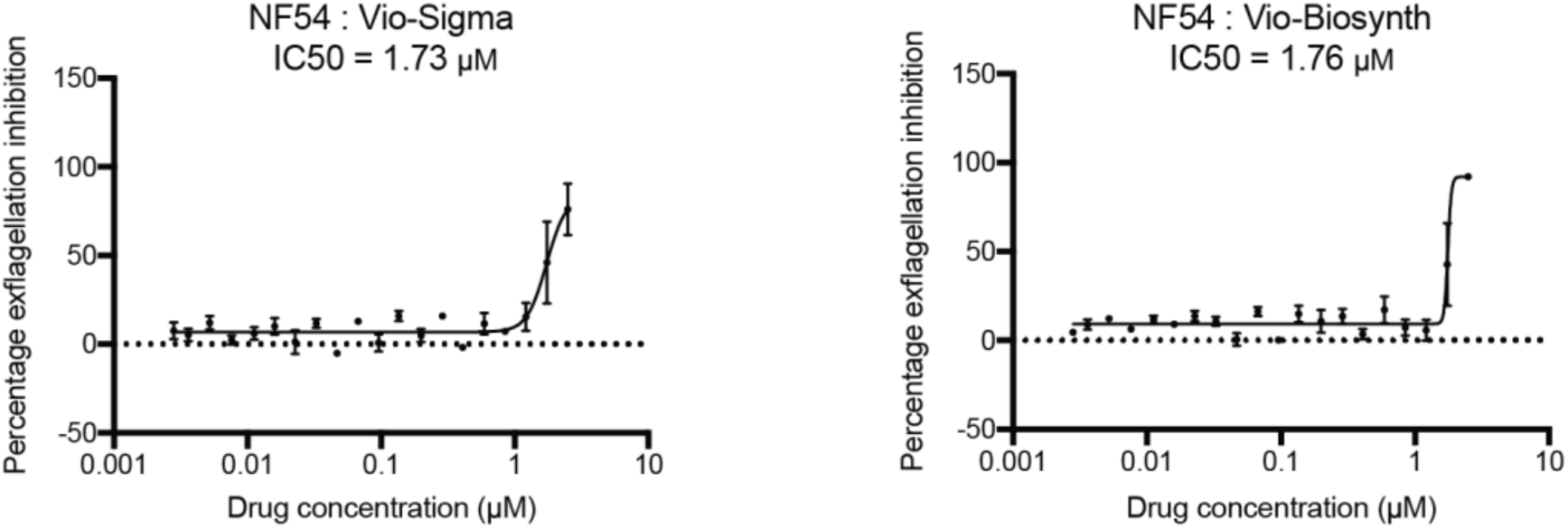
Violacein and biosynthetic violacein inhibit exflagellation with similar IC50 values.

**Fig. S4:**
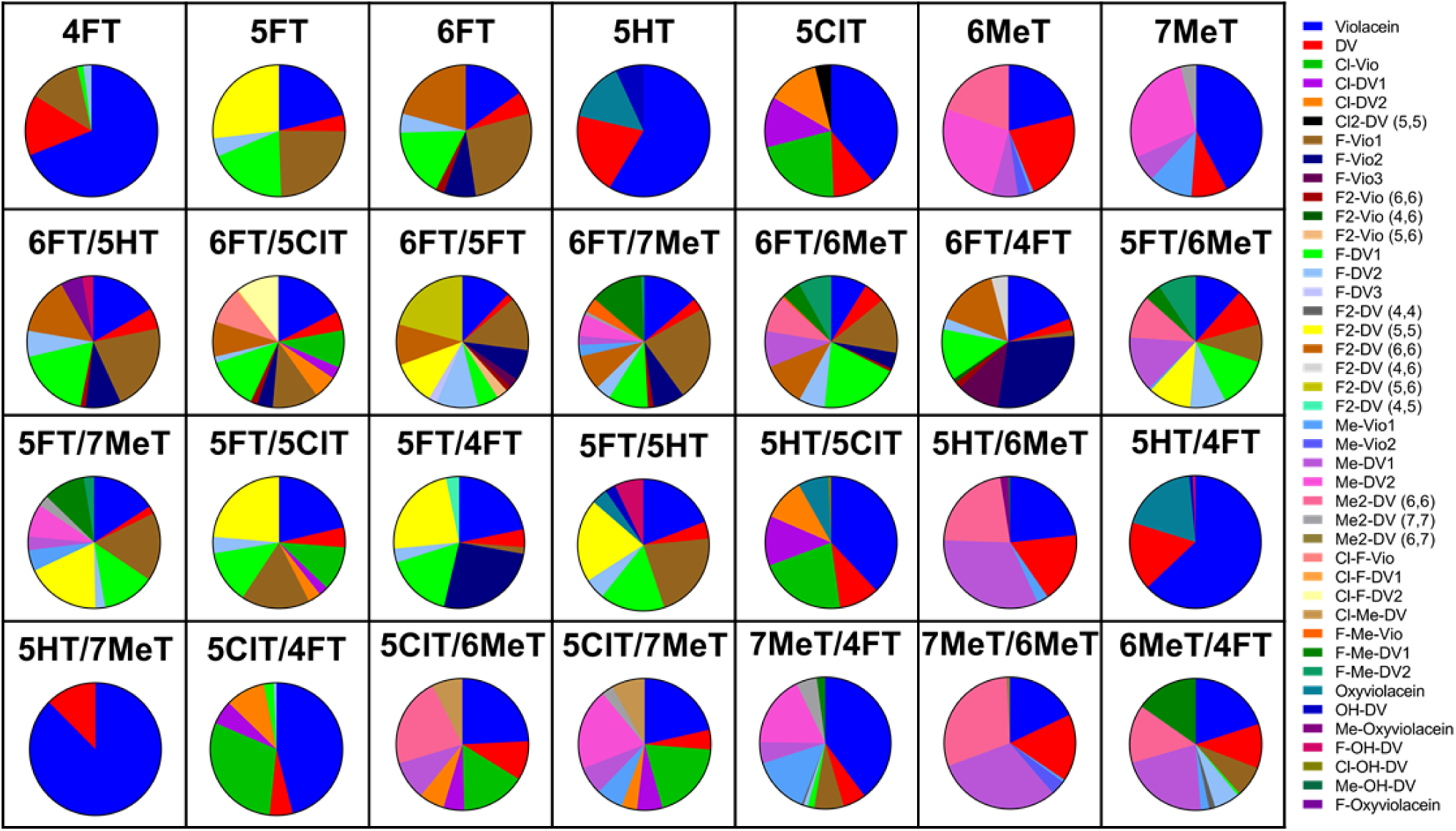
Composition of the violacein derivatives used in the high-throughput single-dose screen. Figure adapted with permission from BioRxiv preprint^24^ (https://www.biorxiv.org/content/10.1101/202523v2).

**Fig. S5:**
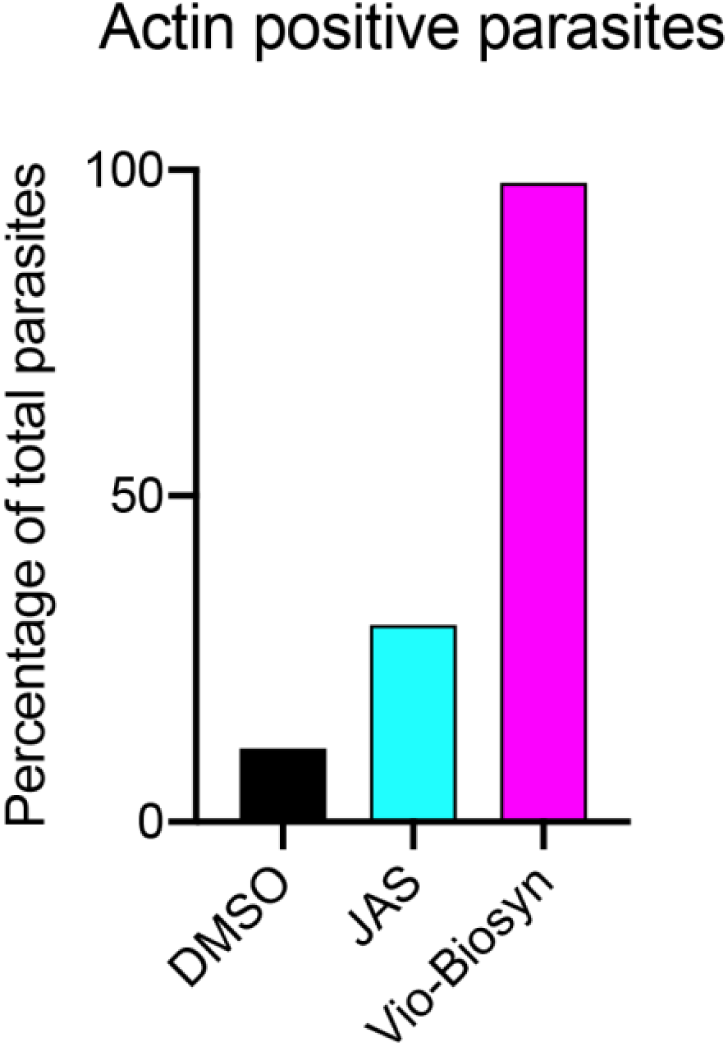
Flow cytometry analysis of fixed 3D7 parasites (100 000 cells sorted). Parasitemia was estimated from an average of the nucleus (DAPI) and cytoplasm (GFP) positive parasites. Actin positive parasites were estimated as a percentage of red-positive cells over total parasitemia.

## REFERENCES

1. WHO | World malaria report 2018. WHO (2019).

2. Saunders, D. L., Vanachayangkul, P. & Lon, C. Dihydroartemisinin–Piperaquine Failure in Cambodia. N. Engl. J. Med. 371, 484–485 (2014).

3. Kanoi, B. N. et al. Antibody profiles to wheat germ cell-free system synthesized Plasmodium falciparum proteins correlate with protection from symptomatic malaria in Uganda. Vaccine (2017). doi:10.1016/j.vaccine.2017.01.001

4. Shaw, P. J. et al. Plasmodium parasites mount an arrest response to dihydroartemisinin, as revealed by whole transcriptome shotgun sequencing (RNA-seq) and microarray study. BMC Genomics 16, 830 (2015).

5. Paloque, L., Ramadani, A. P., Mercereau-Puijalon, O., Augereau, J.-M. & Benoit-Vical, F. Plasmodium falciparum: multifaceted resistance to artemisinins. Malar. J. 15, 149 (2016).

6. Mbengue, A. et al. A molecular mechanism of artemisinin resistance in Plasmodium falciparum malaria. Nature 520, 683–687 (2015).

7. McLaughlin, E. C., Norman, M. W., Ko Ko, T. & Stolt, I. Three-component synthesis of disubstituted 2H-pyrrol-2-ones: preparation of the violacein scaffold. Tetrahedron Lett. 55, 2609–2611 (2014).

8. Petersen, M. T. & Nielsen, T. E. Tandem Ring-Closing Metathesis/Isomerization Reactions for the Total Synthesis of Violacein. Org. Lett. 15, 1986–1989 (2013).

9. Lopes, S. C. P. et al. Violacein Extracted from Chromobacterium violaceum Inhibits Plasmodium Growth In Vitro and In Vivo. Antimicrob. Agents Chemother. 53, 2149–2152 (2009).

10. Pantanella, F. et al. Violacein and biofilm production in Janthinobacterium lividum. J. Appl. Microbiol. 0, 061120055200056-??? (2006).

11. Sneath, P. H. A., Singh, R. B., Whelan, J. P. F. & Edwards, D. FATAL INFECTION BY CHROMOBACTERIUM VIOLACEUM. Lancet 262, 276–277 (1953).

12. Rodrigues, A. L. et al. Systems metabolic engineering of Escherichia coli for production of the antitumor drugs violacein and deoxyviolacein. Metab. Eng. 20, 29–41 (2013).

13. Bilsland, E. et al. Antiplasmodial and trypanocidal activity of violacein and deoxyviolacein produced from synthetic operons. BMC Biotechnol. 18, 22 (2018).

14. Lai, H.-E. et al. A GenoChemetic strategy for derivatization of the violacein natural product scaffold. bioRxiv 202523 (2019). doi:10.1101/202523

15. Rodrigues, A. L. et al. Microbial production of the drugs violacein and deoxyviolacein: analytical development and strain comparison. Biotechnol. Lett. 34, 717–720 (2012).

16. Wang, H. et al. Optimization of culture conditions for violacein production by a new strain of Duganella sp. B2. Biochem. Eng. J. 44, 119–124 (2009).

17. Choi, S. Y., Kim, S., Lyuck, S., Kim, S. B. & Mitchell, R. J. High-level production of violacein by the newly isolated Duganella violaceinigra str. NI28 and its impact on Staphylococcus aureus. Sci. Rep. 5, 15598 (2015).

18. Yeo, A. E. & Rieckmann, K. H. The in vitro antimalarial activity of chloramphenicol against Plasmodium falciparum. Acta Trop. 56, 51–4 (1994).

19. Ruecker, A. et al. A Male and Female Gametocyte Functional Viability Assay To Identify Biologically Relevant Malaria Transmission-Blocking Drugs. Antimicrob. Agents Chemother. 58, 7292–7302 (2014).

20. Malpartida-Cardenas, K. et al. Quantitative and rapid Plasmodium falciparum malaria diagnosis and artemisinin-resistance detection using a CMOS Lab-on-Chip platform. Biosens. Bioelectron. 145, 111678 (2019).

21. Baum, J., Papenfuss, A. T., Baum, B., Speed, T. P. & Cowman, A. F. Regulation of apicomplexan actin-based motility. Nat. Rev. Microbiol. 4, 621–8 (2006).

22. Mbengue, A. et al. A molecular mechanism of artemisinin resistance in Plasmodium falciparum malaria. Nature 520, 683–687 (2015).

23. Miotto, O. et al. Genetic architecture of artemisinin-resistant Plasmodium falciparum. Nat. Genet. 47, 226–234 (2015).

24. Angrisano, F. et al. Spatial Localisation of Actin Filaments across Developmental Stages of the Malaria Parasite. PLoS One 7, e32188 (2012).

25. Johnson, S. et al. Truncated Latrunculins as Actin Inhibitors Targeting Plasmodium falciparum Motility and Host Cell Invasion. J. Med. Chem. 59, (2016).

26. Miguel-Blanco, C. et al. Hundreds of dual-stage antimalarial molecules discovered by a functional gametocyte screen. Nat. Commun. 8, 15160 (2017).

27. Das, S., Lemgruber, L., Tay, C. L., Baum, J. & Meissner, M. Multiple essential functions of Plasmodium falciparum actin-1 during malaria blood-stage development. BMC Biol. 15, 70 (2017).

28. Olshina, M. A., Baumann, H., Willison, K. R. & Baum, J. Plasmodium actin is incompletely folded by heterologous protein-folding machinery and likely requires the native Plasmodium chaperonin complex to enter a mature functional state. FASEB J. (2015). doi:10.1096/fj.15-276618

29. Mok, S. et al. Drug resistance. Population transcriptomics of human malaria parasites reveals the mechanism of artemisinin resistance. Science 347, 431–5 (2015).

